# The population genetics of biological noise

**DOI:** 10.1101/2025.01.11.632402

**Authors:** Daniel M. Weinreich, Tom Sgouros, Yevgeniy Raynes, Hlib Burtsev, Edison Chang, Sanyu Rajakumar, Ignacio G. Bravo, Csenge Petak

**Author notes:** Correspondence and requests for materials should be addressed to Daniel M. Weinreich: Daniel.

## Abstract

Information transmission is intrinsic to life, and noise is intrinsic to information transmission. Biological noise during development is essential for the flexibility and plasticity of individual organisms, but also underlies some diseases. Biological noise during reproduction is the fuel for evolution, including the evolution of therapy resistance in pathogenic microbes and in cancer. Recent technological advances in our ability to characterize many sources of biological noise have demonstrated that its amount is often heritable. Here, we frame the population genetics of loci that influence the amount of any source of biological noise. While analogous theory for heritable changes in mean trait values has been established for nearly a century, to our knowledge this is the first general approach for studying the evolution of heritable changes in their statistical distributions. This represents a critical theoretical contribution to an important and rapidly growing domain of intellectual inquiry. It also sheds light on the hypothesis that natural selection can increase evolvability, and generalizes modifier theory used in the tradition of Feldman and colleagues.

**Author summary:** Biological noise is a fact of life. Genetically identical organisms reared in identical environments invariably exhibit random phenotypic differences. And siblings born of the same parent(s) in the same environment are endowed with inheritances that invariably differ at random. While the specific consequences of biological noise are unpredictable, extraordinary experimental advances now make clear that its amount can be influenced by an organism’s genetics. For example, high- and low-noise promoter, and high- and low-noise DNA polymerase alleles are well known. This raises the question of when and how natural selection favors high- or low-noise alleles. While biological noise is on average deleterious, it can also occasionally induce high fitness phenotypes. Here, we solve a simple analytic model for the fitness difference between noise alleles that captures both these features. Our model predicts the existence of an evolutionary equilibrium in the amount of noise, whose location reflects just four features of an organism’s biology. In light the clinical importance of biological noise, as well as its central role more broadly in both development and evolution, this work provides an urgently needed evolutionary framework for understanding its long-term determinants.

## Introduction

Information transmission is intrinsic to biology: during development, information from an offspring’s inheritance informs its phenotype, and during reproduction, information flows to the next generations’ inheritance (Fig 1A, ref. [1]). But noise is inevitable. During development it results in stochastic fluctuations in biochemical, cellular and organismal phenotypes [2], and during reproduction it causes a newborn’s inheritance to differ from that of its parent(s). Critically, while the consequences of biological noise are random, its amount can be heritably influenced. For example, promoter alleles can differ in gene expression noise (e.g., [3]), and DNA polymerase alleles can differ in their fidelity of genome replication (e.g., [4]).

**Fig 1.**
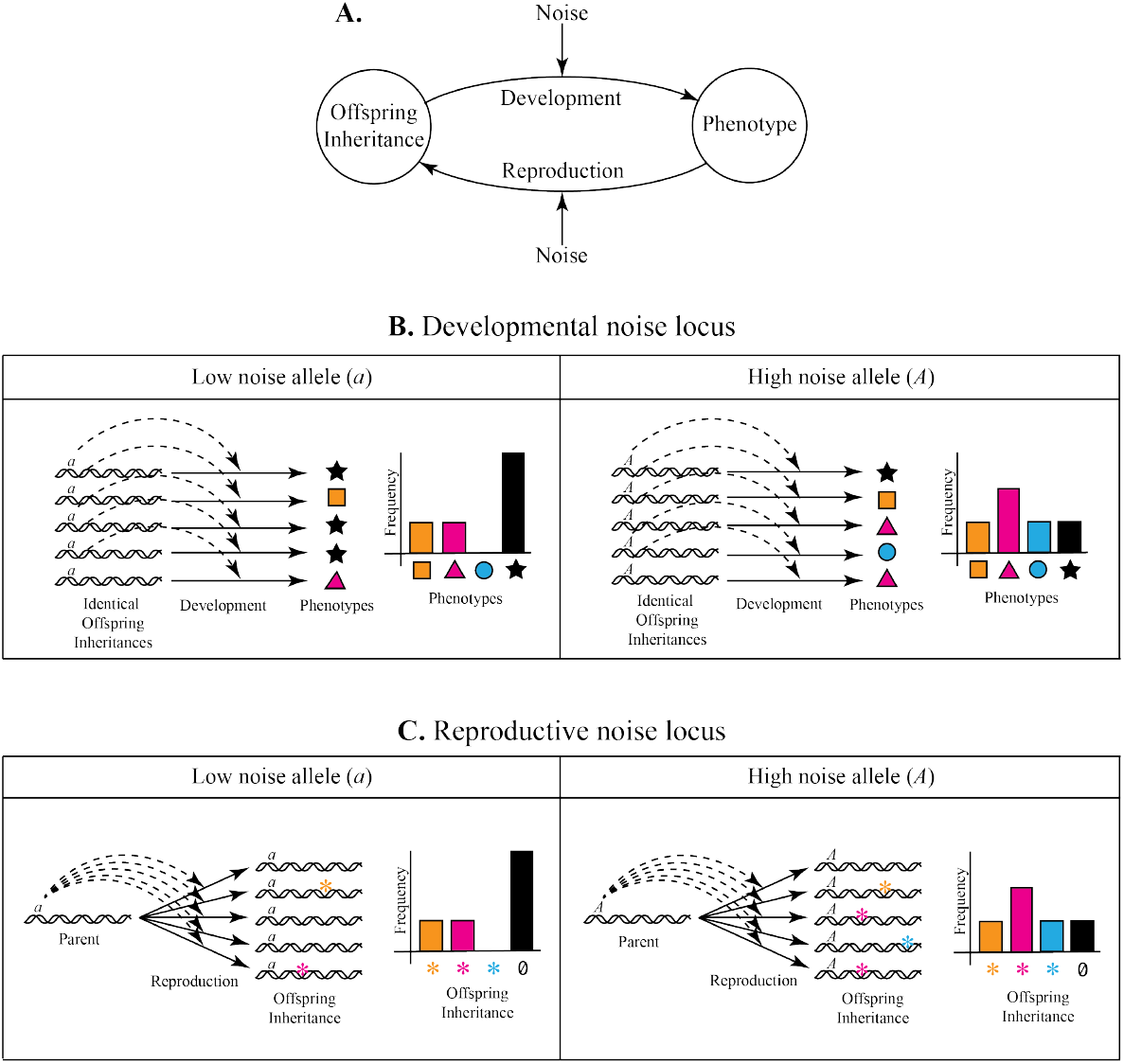
Classes and genetic control of biological noise. **A**. During development, information flows from the offspring’s inheritance to its phenotype, and during reproduction it flows back from the parent to the next generation’s inheritance. This implies that biological noise can emerge during development or during reproduction, correspondingly suggesting two classes of genetic loci that influence the amount of biological noise. **B**. Holding environment constant, low and high developmental noise alleles (designated *a* and *A*) differentially influence the amount of phenotypic variation (represented by colored shapes) among offspring with identical inheritances. **C**. Again holding environment constant, low and high reproductive noise alleles (designated *a* and *A*) differentially influence the amount of variation among offspring inheritances (represented by colored asterisks) from a single parent.

This raises the possibility that alleles that influence the amount of biological noise (hereafter, *noise alleles*) may in some cases be subject to natural selection (e.g., [5, 6]).

We might guess that biological noise is unconditionally deleterious [7, 8], but in fact, it is also occasionally beneficial [9, 10]. Here, we present a simple model for the fitness effects of noise that captures both possibilities. Our model predicts the existence of an equilibrium amount of biological noise, the location of which depends on four factors: the rate of environmental change, the timescale of association between noise allele and induced phenotype, the form of selection, and population size. We argue that this result may be robust to model specifics, and conclude by connecting it to previous art and to future work.

## Results

The distinction between developmental and reproductive noise (Fig 1A) motivates us to distinguish between *developmental* and *reproductive noise alleles*, which reside respectively at *developmental* and *reproductive noise loci*. Fig 1B schematically illustrates how developmental noise alleles influence phenotypic variation among offspring with identical inheritance. And Fig 1C shows how reproductive noise alleles affect random variation in inheritance among offspring of the same parent. (We assume throughout that variation in inheritance causes phenotypic variation.) Table 1 presents many mechanisms for both classes of biological noise whose intensity is under genetic control.

**Table 1:**
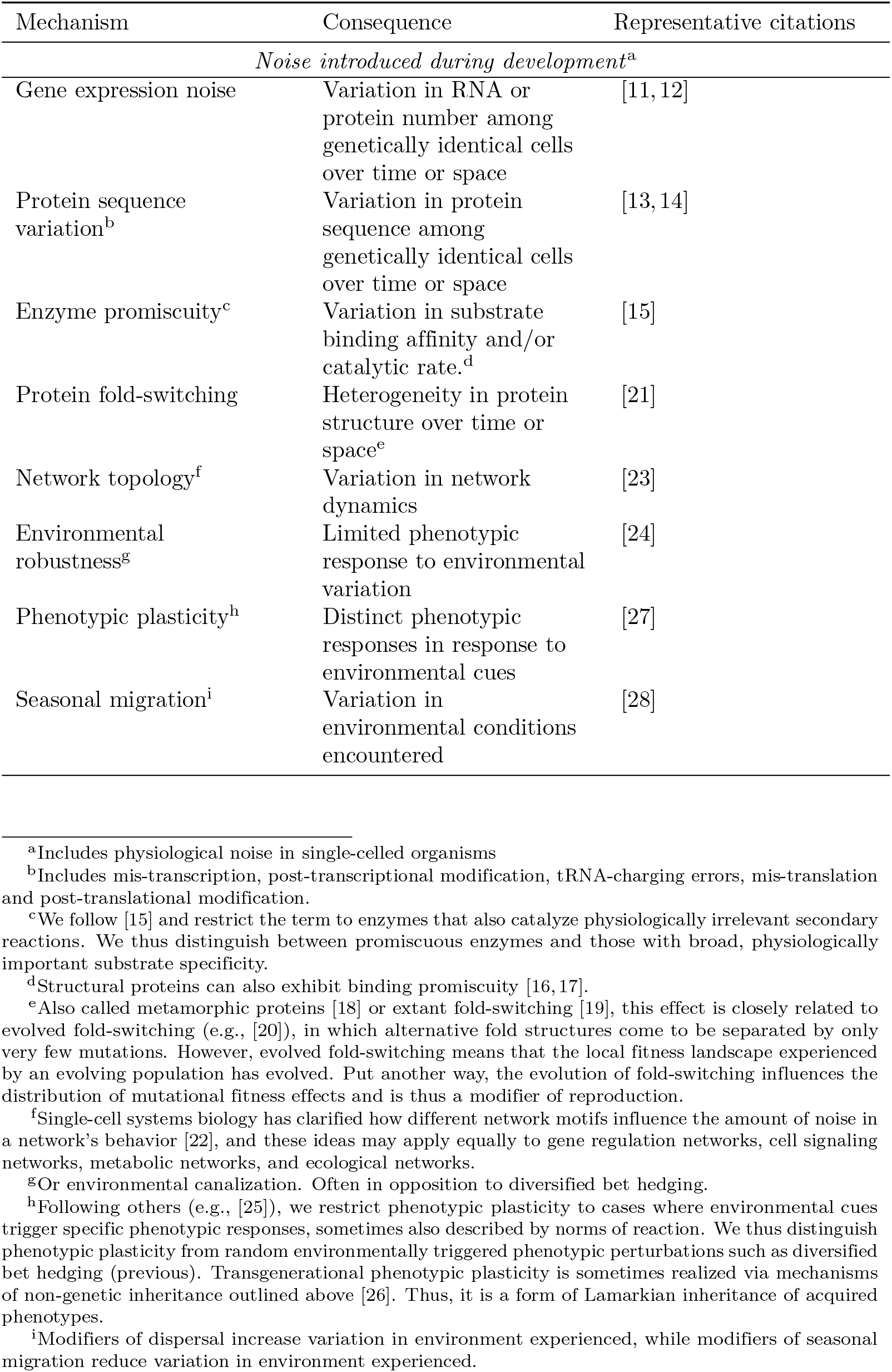
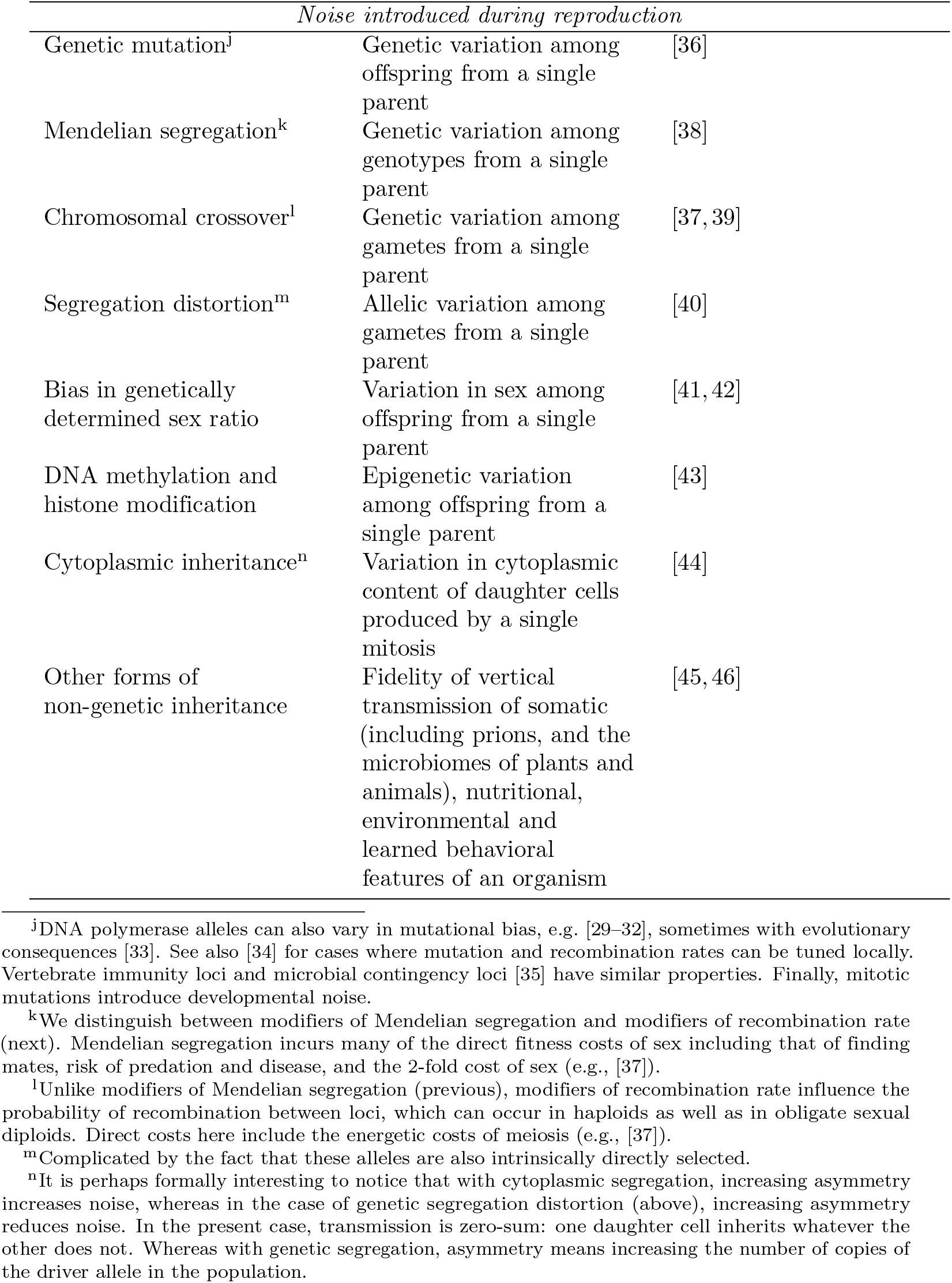
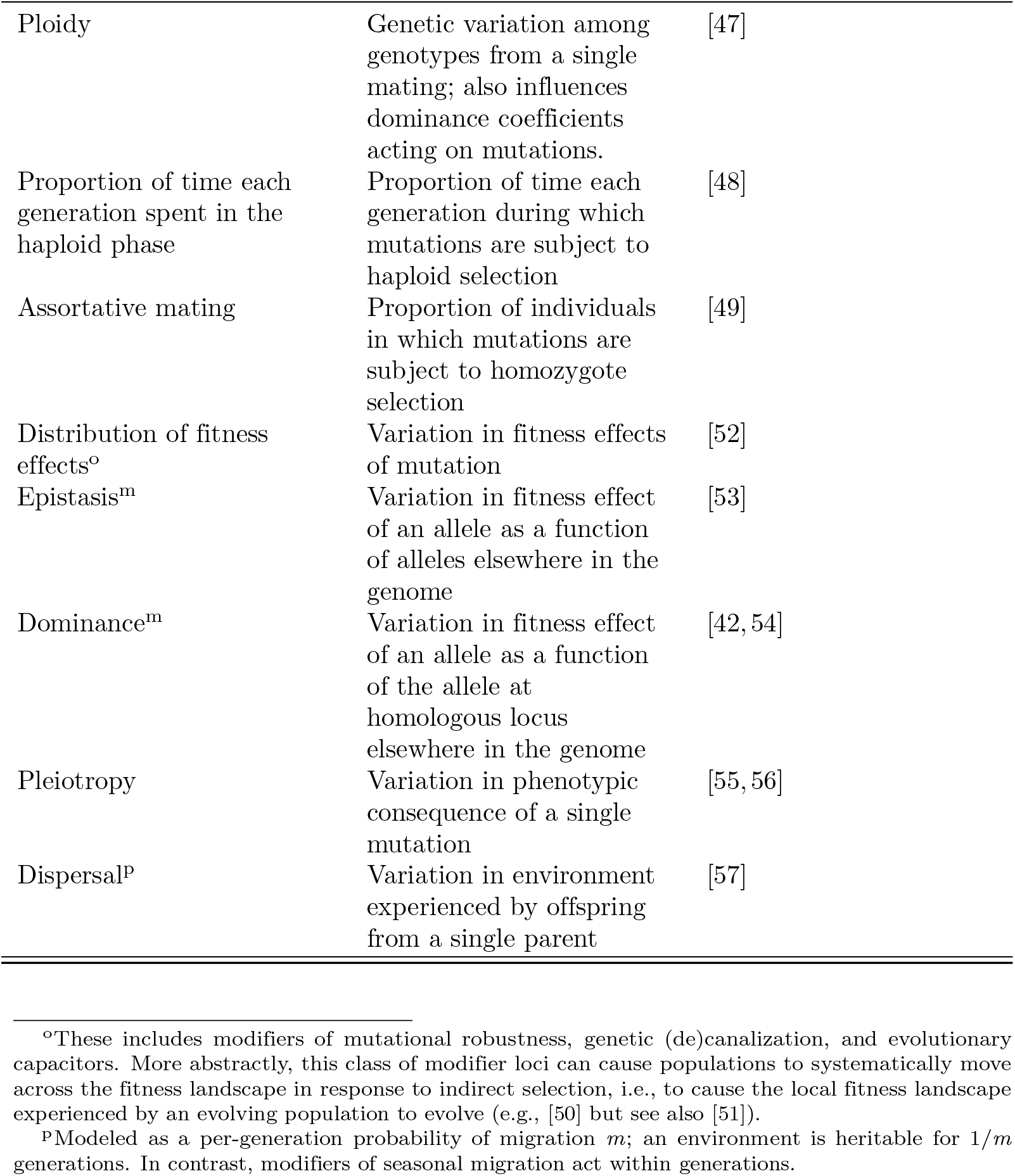
Mechanisms and consequences of biological noise whose intensity may be under genetic control.

| Mechanism | Consequence | Representative citations |
| --- | --- | --- |
| *Noise introduced during development<sup>a</sup>* |  |  |
| Gene expression noise | Variation in RNA or protein number among genetically identical cells over time or space | [11, 12] |
| Protein sequence variation<sup>b</sup> | Variation in protein sequence among genetically identical cells over time or space | [13, 14] |
| Enzyme promiscuity<sup>c</sup> | Variation in substrate binding affinity and/or catalytic rate.<sup>d</sup> | [15] |
| Protein fold-switching | Heterogeneity in protein structure over time or space<sup>e</sup> | [21] |
| Network topology<sup>f</sup> | Variation in network dynamics | [23] |
| Environmental robustness<sup>g</sup> | Limited phenotypic response to environmental variation | [24] |
| Phenotypic plasticity<sup>h</sup> | Distinct phenotypic responses in response to environmental cues | [27] |
| Seasonal migration<sup>i</sup> | Variation in environmental conditions encountered | [28] |
| *Noise introduced during reproduction* |  |  |
| Genetic mutation<sup>j</sup> | Genetic variation among offspring from a single parent | [36] |
| Mendelian segregation<sup>k</sup> | Genetic variation among genotypes from a single parent | [38] |
| Chromosomal crossover<sup>l</sup> | Genetic variation among gametes from a single parent | [37, 39] |
| Segregation distortion<sup>m</sup> | Allelic variation among gametes from a single parent | [40] |
| Bias in genetically determined sex ratio | Variation in sex among offspring from a single parent | [41, 42] |
| DNA methylation and histone modification | Epigenetic variation among offspring from a single parent | [43] |
| Cytoplasmic inheritance<sup>n</sup> | Variation in cytoplasmic content of daughter cells produced by a single mitosis | [44] |
| Other forms of non-genetic inheritance | Fidelity of vertical transmission of somatic (including prions, and the microbiomes of plants and animals), nutritional, environmental and learned behavioral features of an organism | [45, 46] |
| Ploidy | Genetic variation among genotypes from a single mating; also influences dominance coefficients acting on mutations. | [47] |
| Proportion of time each generation spent in the haploid phase | Proportion of time each generation during which mutations are subject to haploid selection | [48] |
| Assortative mating | Proportion of individuals in which mutations are subject to homozygote selection | [49] |
| Distribution of fitness effects<sup>o</sup> | Variation in fitness effects of mutation | [52] |
| Epistasis<sup>m</sup> | Variation in fitness effect of an allele as a function of alleles elsewhere in the genome | [53] |
| Dominance<sup>m</sup> | Variation in fitness effect of an allele as a function of the allele at homologous locus elsewhere in the genome | [42, 54] |
| Pleiotropy | Variation in phenotypic consequence of a single mutation | [55, 56] |
| Dispersal<sup>p</sup> | Variation in environment experienced by offspring from a single parent | [57] |

<sup>o</sup>These includes modifiers of mutational robustness, genetic (de)canalization, and evolutionary capacitors. More abstractly, this class of modifier loci can cause populations to systematically move across the fitness landscape in response to indirect selection, i.e., to cause the local fitness landscape experienced by an evolving population to evolve (e.g., [50] but see also [51]).
<sup>p</sup>Modeled as a per-generation probability of migration $m$ ; an environment is heritable for $1/m$ generations. In contrast, modifiers of seasonal migration act within generations.

### Modeling the fitness effects of biological noise

Since the phenotypic consequence of biological noise is random, we introduce the stochastic model shown in Fig 2A. We assume a one-dimensional, normally distributed phenotype *z*:

**Fig 2.**
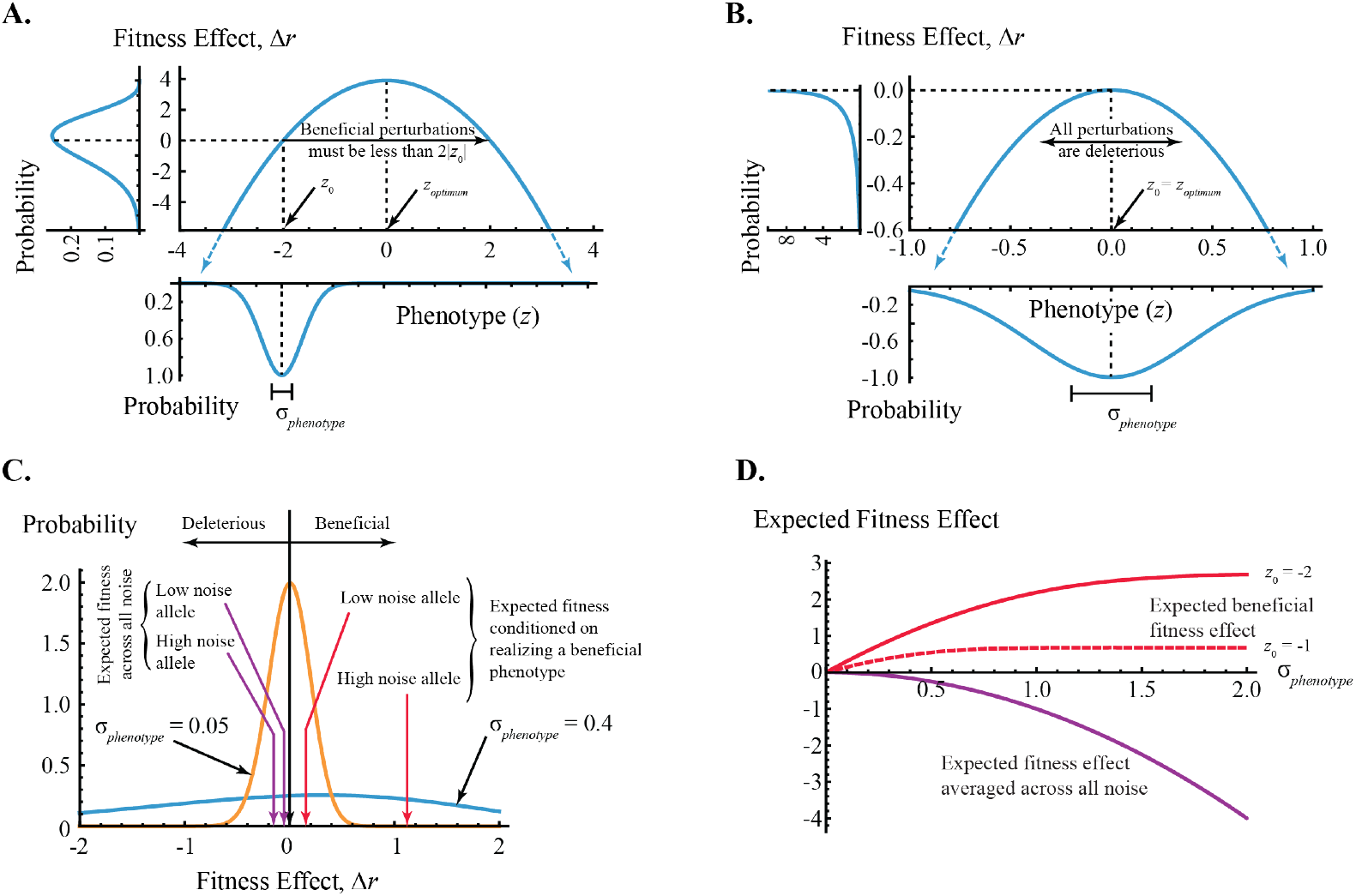
Fitness effects of phenotypic noise. **A**. Model introduced in text. Lower plot: Normally distributed phenotypic noise (Eq. 1) with expectation *z*_0_ = 𝔼[*z*] = -2 and phenotypic noise *σ*_*phenotype*_ = 0.4. Center plot: Quadratic phenotype-to-fitness function (Eq. 2). Without loss of generality we set the phenotypic fitness optimum *z*_*optimum*_ = 0. Left plot: resulting probability density function of fitness effects Δ*r* (Eq. 3). **B**. All phenotypic noise is deleterious if an organism’s expected phenotype is at the fitness optimum (*z*_0_ = 𝔼[*z*] = *z*_*optimum*_ = 0). All other parameters as in panel A, though the x-axis has been rescaled for emphasis. **C**. The probability density functions of fitness effects induced by high (blue, *σ*_*phenotype*_ = 0.4, as in panel A) and low (orange, *σ*_*phenotype*_ = 0.05) noise alleles. All other parameters as in panel A, though the x-axis has been rescaled for emphasis. In expectation, noise is deleterious. This can be seen by comparing the two purple arrows with the vertical black arrow. But as noise goes up, so does the expected fitness effect of beneficial phenotypes. This can be seen by comparing two red arrows with the vertical black arrow. **D**. Expected fitness averaged over all phenotypes (purple, given by Eq. 4; this is independent of expected phenotype) and conditioned on beneficial phenotypes (red, Eq. 5; shown for two values of expected phenotype *z*_0_) as a function of phenotypic noise (*σ*_*phenotype*_). In the latter case, solid line correspond to the case when expected phenotype *z*_0_ = -2 (as also shown in panels A and C) while dashed line show result when expected phenotype *z*_0_ = -1.

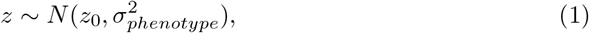

where *σ*_*phenotype*_ is our proxy for the amount of phenotypic noise and *z*_0_ is the organism’s expected phenotype averaged over noise (equivalently, *z*_0_ = 𝔼[*z*]). The form of Eq. 1 is shown at the bottom of Fig 2A.

Following others (e.g., [58]), we next assume that fitness (*r*) drops quadratically in *z* from an optimal phenotype (*z*_*optimum*_), written *r* (*z*) = *r*_*optimum*_ *−* (*z*_*optimum*_ *− z*)^2^. Without loss of generality, we set the optimal phenotype *z*_*optimum*_ = 0. Thus, the fitness effect of a phenotypic perturbation to *z* in an organism with expected phenotype *z*_0_ is

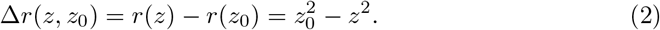

These ideas are illustrated in the center of Fig 2A. Note that our model assumes that biological noise influences only a single phenotype that is sampled exactly once each generation. We address both assumptions in the Discussion.

Finally, the change-of-value technique (Methods) yields an expression for the probability density function of fitness effects (Eq. 3) for given expected phenotype (*z*_0_) and phenotypic noise (*σ*_*phenotype*_). Typical results are shown at the left of Fig 2A and B,and in 2C.

### Biological noise is deleterious on average

In our model, all phenotypic perturbations larger than 2 |*z*_0_| are deleterious (Fig 2A), and since we assume that noise is symmetric in *z*, only half the remainder are beneficial. Moreover, for a given phenotypic perturbation less than 2 |*z*_0_|, deleterious outcomes will be more deleterious than beneficial outcomes are beneficial. This effect, called Jensen’s inequality [59], follows because fitness effect Δ*r* is concave down. (Algebraically, Jensen’s inequality reads 𝔼[Δ*r* (*z,z*_0_)] *<* Δ*r* (𝔼[*z*],*z*_0_).) Thus on expectation, fitness goes down as noise goes up, an effect confirmed quantitatively by Eq. 4 (Methods).

The limiting case is an organism at the phenotypic fitness optimum (*z*_0_ = 𝔼[*z*] = *z*_*optimum*_ = 0). Here, all perturbations are deleterious (Fig 2B). Indeed, even in populations not quite at the phenotypic optimum but only at equilibrium between natural selection, mutation, recombination, migration and demographic stochasticity within lineages (called *random genetic drift*), noise due to mutation, migration, or recombination is deleterious [60]. This result is called the *reduction principle*, and we speculate that it generalizes to all sources of biological noise.

### Noise can sometimes be beneficial

But while the fitness effect of noise averaged across all phenotypes goes down as noise goes up, in the more general case of an organism not at the phenotypic fitness optimum, increased noise also increases access to high fitness phenotypes. This can be seen by comparing the low noise and high noise probability density functions in Fig 2C. We quantify this second effect by also computing the expected fitness effects conditioned on the realization of a beneficial phenotype (Eq. 5, Methods). As suggested at the outset, the tension between these *unconditioned* and *conditioned fitness expectations* (algebraically, 𝔼[Δ*r*(*z, z*_0_)] and 𝔼[Δ*r*(*z, z*_0_) | Δ*r*(*z, z*_0_) *>* 0)], respectively) is critical to understanding the evolutionary fate of noise alleles.

Fig 2D illustrates the dependence of both expectations on *σ*_*phenotype*_ under our model. The expected fitness across all phenotypes (purple) goes down without limit as noise goes up (as per Jensen’s inequality). Interestingly, this effect is independent of expected phenotype *z*_0_ (Eq. 4). In contrast, the conditioned expectation (red) goes up before approaching a plateau. That plateau goes down as the expected phenotype approaches the optimum.

### An equilibrium amount of biological noise

The two features of our model represented by the changing slopes of the purple and red lines in Fig 2D suggest the existence of an equilibrium amount of biological noise. From there, higher noise alleles will be selectively disfavored because their now more modest conditioned fitness advantage is more than offset by their greater unconditioned fitness cost. And at the same time, lower noise alleles will also be selectively disfavored, because their inability to capitalize on anomalously high fitness phenotypic realization is not offset by their reduced unconditioned fitness cost. We identify four determinants for the location of this equilibrium: the rate of environmental change, the timescale of association between noise alleles and the phenotypic perturbations they induce, the form of natural selection, and population size.

### Rate of environmental change

Under our model, the scope for fitness improvement induced by biological noise goes down as an organism’s expected phenotype *z*_0_ approaches the phenotypic fitness optimum *z*_*optimum*_ = 0 (compare solid and dashed red lines in Fig 2D.) Now relaxing the assumption that the environment (and hence, the fitness function) is invariant in time, we imagine an evolving population as lagging behind its moving phenotypic fitness optimum in phenotypic space by an amount that goes up with the rate of environmental change. Thus, we predict that the equilibrium amount of biological noise similarly goes up with the rate of environmental change.

### Timescales of phenotypic association

Indirect selection (Box 1) can only distinguish between segregating noise alleles in a population via whatever directly selected phenotypic differences they happen to have recently induced in their carriers. Thus, indirect selection’s influence on the amount of biological noise in a population must somehow depend on the timescale of association (in organismal generations) between noise alleles and their resulting phenotypic perturbations. Put another way, indirect selection’s efficacy will depend on the correlation between parent and offspring phenotype.

One limit is represented by alleles that influence mutation rate in non-recombining organisms. These remain permanently associated with whatever phenotypic perturbations they cause via mutations elsewhere in the genome. Thus here, phenotypic correlation between parent and offspring is maximized, allowing indirect selection to distinguish between competing noise alleles until one reaches fixation and the other is lost. This is why high noise mutation rate alleles (or *mutators*) often reach fixation in asexual microbial populations evolving in novel lab environments [73–78]. This also explains the converse observation: low noise mutation rate alleles (or *antimutators*) often reach fixation in asexual populations evolving in an environment to which they are already well adapted ([74, 77, 79–81], though see [82]). (Note that these two results are consistent with our prediction that the equilibrium amount of biological noise should be positively correlated with the pace of environmental change.)

At the other limit, consider a noise allele for which parent and offspring phenotypes are entirely uncorrelated. In this case indirect selection always disfavors high noise alleles [83]. This follows because while still rare, the risk that the high noise allele is lost in the next generation as a consequence of sampling a deleterious phenotype is of much greater evolutionary significance than is the (one-generation) benefit of increased frequency as a consequence of its sampling a beneficial one.

In between these limits, the probability of success for a new high noise allele goes up with the timescale of association with its induced phenotypes [84], since this allows indirect selection to “see” its occasionally beneficial phenotypic consequences for more generations. Thus, all else equal, we predict that the equilibrium amount of biological noise similarly goes up with the timescale of association.

This timescale is determined by two biological processes: the rate of dissociation between noise allele and induced phenotype (e.g., via recombination in the case of mutator alleles [42]), and the lability of the induced phenotype itself. To illustrate this latter process, consider a locus in a single-celled organism that influences the variance in cellular concentration of some biomolecule. Now imagine an individual in which a high noise allele has induced a selectively favorable, anomalously high (or low) concentration of that molecule. While subsequent cell divisions will dilute such perturbations back toward its expectation, that decay will be roughly exponential, meaning that indirect selection will favor the high noise allele for a few cell generations. Similarly, transgenerational phenotypic plasticity in multicellular organisms often persists for several generations after the environmental cue (reviewed in [26]), again offering indirect selection a reliable signal with which to distinguish between competing noise alleles.

#### Box 1: Direct and indirect selection

Classical population genetic models of natural selection disregard biological noise, instead assuming that any given allele has the identical phenotypic effect on every otherwise identical organism in which it appears. In this case, the fate of an allele can be modeled by recursively applying that effect to every descendant carrier in which it may appear [61]. Extending usage of others (e.g., [36, 42, 62]), we say that such alleles are under *direct selection*.

In contrast, noise alleles induce stochastic fitness perturbations in their carriers as a consequence of stochastic perturbations in phenotype. Thus, natural selection can influence the frequency of a noise allele via direct selection on whatever phenotype it happens to have most recently induced in its carrier. Following others (e.g., [36, 42, 62]), we say that such alleles are under *indirect selection*. (This is also sometimes called *second order selection*, e.g., [63–65].) Critically, indirect selection allows for the possibility that two copies of the same noise allele may be regarded differently by natural selection, substantially complicating the problem.

Importantly, noise alleles can also be directly selected. For example, fitness in some RNA viruses depends in part on genome replication rate [66, 67]. Thus, an RNA polymerase allele that is faster because it lacks a proofreading activity can accelerate genome replication, thereby uniformly increasing fitness in all carriers. Such direct selection is independent of any indirect selection mediated by the phenotypic consequences of whatever additional mutations the error-prone polymerase introduces. Similarly, on the premise that misfolded proteins are metabolically or physiologically costly, alleles that reduce chaperone protein expression will be uniformly deleterious in all carriers. Again, this direct effect is independent of any indirect selection acting on the phenotypic consequences of a decrease in the proportion of mis-folded proteins.

The distinction between direct and indirect selection goes back to Fisher [68]. Following others (e.g., [62, 69]), here we simplify by assuming that noise alleles only experience indirect selection. Algebraically, this means we assume that cosegregating noise alleles only differ in *σ*_phenotype_ and not in *z*_0_. The interested reader is directed to [42] and [52], where this assumption is explicitly relaxed. See also [70], [71], and [72], which suggest that this simplification may sometimes be biologically untenable.

Finally, recall that the number of generations from appearance to fixation of a directly selected beneficial mutation is roughly the reciprocal of its fitness advantage [85]. Thus, indirect selection’s capacity to favor high noise alleles is maximized as soon as phenotypic association times are at least that long.

### The form of selection

Results thus far have assumed fitness-proportional selection, meaning that the probability that an organism contributes to the next generation is proportional to its fitness in the current one. But other effects are possible. For example, when males compete for reproductive access to females, all individuals that exceed some phenotypic threshold can have an equal probability of contributing to the next generation. This is called truncation selection, and it emerges in many biological situations (reviewed in [86, 87]).

Interestingly, [86] showed that truncation selection can cause indirect selection to favor high noise alleles even if phenotypes are resampled every generation. In this case, individuals contributing to the next generation will include carriers of high noise alleles that happen by good luck to have sampled a phenotype above the threshold, even if their expected phenotype is below that value. Put another way, truncation selection reduces the fitness penalty induced by high noise alleles. Indeed, even less extreme departures from fitness-proportional selection can be sufficient to favor high noise alleles in the absence of any persistent association with induced phenotype [86]. Consequently, we predict that the amount of noise expected at the selective equilibrium will go up as selection goes from fitness-proportional toward truncation.

### Population size

For a population at its phenotypic fitness optimum, all noise is deleterious (Fig 2B). And since the strength of random genetic drift scales inversely with population size [41, 88], we predict that in such populations, the equilibrium amount of noise should go down as population size goes up. This is Lynch’s *drift barrier* hypothesis, [89]), which is consistent with the pattern observed for mutation rate across cellular life (*r* ^2^*≈* 0.8, [90]). The analogous signal is weaker for single-point transcription [91, 92] and splicing [93] error rates. Though we might not be surprised: the consequence of mutational noise are transmitted to all offspring, whereas transcription and splicing noise only affect the products of individual mRNA molecules.

Importantly, the picture is more complicated in the general case of populations not at the phenotypic fitness optimum, because some stochastic phenotypic fluctuations are now beneficial (Fig 2A, C). The strength of random genetic drift still goes down as population size goes up, and assuming that most phenotypic perturbations are deleterious, indirect selection against high noise alleles will be the first effect to overcome drift [94]. Thus in populations of moderate size, we still predict a drift barrier effect (i.e., selection against the high noise allele). But as population size continues to increase, the strength of indirect selection in favor of high noise alleles will eventually come to exceed both the strength of random genetic drift and indirect selection against them [94]. Thus, we predict that at above some critical population size, the equilibrium amount of noise will go up with population size.

This non-monotonicity in the direction of selection acting on noise alleles is called *sign inversion* [95]. It emerges from the double-edged nature of biological noise and has been observed in several systems [47, 95–101]. We speculate all noise alleles will exhibit sign inversion in populations not at their phenotypic fitness optimum.

## Discussion

Our treatment of the population genetics of biological noise is predicated on an intrinsic tension between the beneficial phenotypes that noise occasionally induces and its much more common, deleterious consequences. We have argued that for any biological mechanism in which the amount of noise is under genetic control, there will exist an equilibrium amount at which these opposing effects are precisely balanced. This result depends on the distinction between direct and indirect selection (Box 1), which is also central to two other active domains of intellectual inquiry.

### Prior art

First, indirect selection is central to theory that explores the evolution of *evolvability*, a population’s capacity to generate heritably adaptive phenotypic variants [70, 102–104]. More specifically, the distinction between evolvability-as-byproduct and evolvability-as-adaptation [70, 105] is equivalent to the distinction between direct and indirect selection acting on evolvability. Critically however, evolvability-as-adaptation arguments risk teleological critiques, since an evolving population cannot “anticipate” what phenotypes are likely to be beneficial in the future (e.g., [27, 70, 106, 107]).

Reframing the problem of indirect selection on evolvability in terms of biological noise appears to resolve this concern, since noise is inherently a double-edged sword. As we have shown, indirect selection increases biological noise only when increasing the population’s capacity to generate adaptive phenotypic variants (i.e., its evolvability) outweigh the concomitant cost of generating deleterious variants. Otherwise, indirect selection will reduce biological noise because this reduction in the population’s capacity to generate deleterious phenotypes (what might be called its *antievolvability*) now outweighs the possibility of generating adaptive phenotypes.

Indirect selection is also fundamental to a body of evolutionary literature called *modifier theory*. That work begins from the premise that population genetic parameter values such as mutation, recombination and migration rates are genetically encoded by mutable *modifier alleles*, which thus can evolve. Modifier theory dates to the 1960s [63, 69, 108], and the field remains quite active (e.g., [60, 96, 99, 109]).

We believe that our approach represents a generalization of modifier theory. Nearly all work in modifier theory has focused on the evolution of parameter values that influence reproductive noise (Table 1; though see [96] for one exception). Correspondingly, modifier theory recognizes the disruptive effect of genetic recombination on indirect selection [42]. Our focus on the timescales of association between noise allele and induced phenotype is an abstraction of this effect, suggesting that results from modifier theory may also apply to developmental noise alleles. Moreover, our focus on mechanisms of noise rather than on parameter values in population genetic models hints that a deeper understanding of these phenomena may be possible. These ideas are being explored in a forthcoming contribution.

### Future work

Importantly, our model makes several strong assumptions. For example, in contrast to Fig 2, noise alleles can have multidimensional phenotypic consequences, such as when they influence the pleiotropic effects of mutations [55, 56, 110, 111]. Moreover, regardless of dimensionality, we have limited intuition for the distribution of fitness effects of phenotypic noise. And experimental access to these distributions is difficult for most of the mechanisms in Table 1, though surveys of the distribution of mutational fitness effects are one exception (e.g., [112–114]).

Nevertheless, our prediction of an equilibrium amount of biological noise may be quite robust to model specifics. First, the existence of a nearby optimum on the fitness function appears to us to be sufficient for the conditioned, beneficial effect of noise to approach a plateau as noise goes up. Second, recent work on mutational interactions reveals that as fitness goes up, the advantage of beneficial mutations often decline while the costs of deleterious mutations go up (reviewed in [115, 116]). By Jensen’s inequality, such negative curvature in the phenotype-to-fitness function is sufficient to cause the unconditioned fitness of noise alleles to go down with noise. It remains to be seen whether the fitness function acting on other forms of biological noise also exhibit negative curvature, although there are principled reasons to suppose that they may [8, 117–119]).

Another critical issue outside our treatment is the interplay between the rate of environmental variation and that of organismal development. Our model assumes that organisms sample phenotypes once in their lifetime. However, physiology can change depending on the environment, and development is not an instantaneous process. So noise may influence phenotype (and thus fitness) more than once during an individual’s lifetime. This is perhaps especially important in long-lived organisms with distinct developmental phases. Moreover, the distributions from which these perturbations are sampled can themselves vary over an organism’s lifetime (e.g., [120, 121]). Similarly, in multicellular organisms developmental noise can be sampled independently across cells [122] or cell types [123]. It appears plausible that if the environment varies substantially within a generation, changes in the phenotype may magnify the evolutionary importance of developmental noise alleles, but much work remains to be done in this area.

Finally, we have only presented a model for the distribution of fitness effects of biological noise. A complete treatment of the resulting population genetics remain to be solved. Perhaps of greatest importance, we require an analytic expression for the equilibrium amount of biological noise as a function of the four determinants introduced above. Relatedly, it will be important to gain insights into the probability of (and time to) fixation for new noise alleles. As noted (Box 1), these are challenging questions, although some results are known (e.g., [83, 95, 98, 124]).

The population genetics of alleles that influence mean trait values have been well established for almost a century. Here we propose an original approach for framing the population genetics of alleles that influence the amount of biological noise around those means. This work is extremely timely because technological advances are revolutionizing empirical access to this phenomenon at the cellular, subcellular and molecular levels (Table 1). Biological noise also has important implications for human disease (e.g., [122, 125]) and the evolution of therapy resistance of pathogenic microbes and in cancer. We are optimistic that the ideas introduced here will soon lead to a much deeper understanding of the forces responsible for observed levels of biological noise in nature.

## Methods

If phenotype *z ∼N* (*z*_0_, *σ*^2^) (Eq. 1) and fitness effect 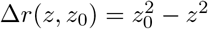 (Eq. 2), the change-of-value technique (shown here) gives the probability density function for fitness effects Δ*r*(*z*):

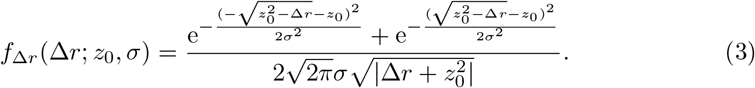

This in turn yields the expected fitness effect over all phenotypic perturbations:

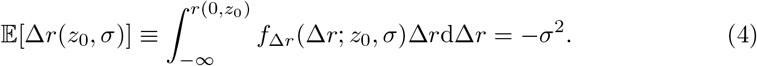

(Recall that without loss of generality we set *z*_optimum_ = 0, making *r* (0,*z*_0_) the appropriate upper limit of integration.)

Finally, the expected fitness effect conditioned on a beneficial phenotype reads

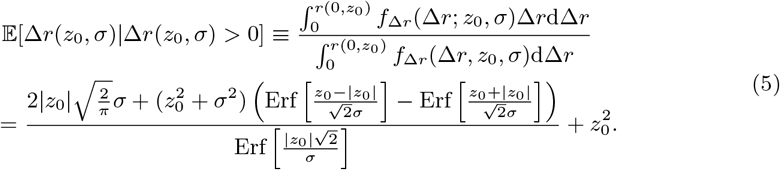

All calculations were performed in Mathematica, as was the production of Fig 2. That code is available [here](https://doi.org/10.5281/zenodo.15591471) under GNU General Public License (GPLv3), and can be examined with the free [Wolfram Player application](https://www.wolfram.com/player/). Eqs. 3-5 match simulation results developed in MatLab (not shown; available on request from daniel).

## Acknowledgments

This work emerged from a graduate seminar at Brown University that DMW taught during the spring semester of 2020, and we thank Nigel Anderson, Alan Denadel, Gabriella Ferra, Lourdes Gomez, Jiaying Lai, Robert Lamb, Miles, David Morgan, Chibuikem Nwizu, Maya Weissman and Hannah Weller for early contributions. Guillaume Achaz, Christina Burch and Ricardo Azevedo provided several trenchant suggestions late in the writing. DMW is grateful for financial support from the Brown University Faculty Research Fund during his Fall 2024 sabbatical in Montpellier, France, during which much of this work was completed. He also acknowledges space and intellectual support during that visit from the *Équipe Génétique et Écologie Évolutive* at the *Centre d’Écologie Fonctionnelle et Évolutive* in Montpellier, and by members of the *Maladies Infectieuses et Vecteurs: Écologie, Génétique, Évolution et Contrôle* laboratory in Montpellier. IGB was supported by the European Union’s Horizon 2020 Research and Innovation Program (ERC-CoG-647916).

## Author Contributions

DMW conceived the idea for this manuscript, DMW, CP, TS, YR and IGB developed the conceptual framework, DMW wrote and solved the model, and DMW, CP, TS and IGB wrote the manuscript. DMW, CP, HB, EC, and SR conducted the literature review.

